# Quantitative benzimidazole resistance and fitness effects of parasitic nematode beta-tubulin alleles

**DOI:** 10.1101/2020.07.07.191866

**Authors:** Clayton M. Dilks, Steffen R. Hahnel, Qicong Sheng, Lijiang Long, Patrick T. McGrath, Erik C. Andersen

## Abstract

Infections by parasitic nematodes inflict a huge burden on the health of humans and livestock throughout the world. Anthelmintic drugs are the first line of defense against these infections. Unfortunately, resistance to these drugs is rampant and continues to spread. To improve treatment strategies, we must understand the genetics and molecular mechanisms that underlie resistance. Studies of the fungus *Aspergillus nidulans* and the free-living nematode *Caenorhabditis elegans* discovered that a beta-tubulin gene is mutated in benzimidazole (BZ) resistant strains. In parasitic nematode populations, three canonical beta-tubulin alleles, F200Y, E198A, and F167Y, have long been correlated with resistance. Additionally, improvements in sequencing technologies have identified new alleles - E198V, E198L, E198K, E198I, and E198Stop - also correlated with BZ resistance. However, none of these alleles have been proven to cause resistance. To empirically demonstrate this point, we independently introduced the three canonical alleles as well as two of the newly identified alleles, E198V and E198L, into the BZ susceptible *C. elegans* N2 genetic background. These genome-edited strains were exposed to both albendazole and fenbendazole to quantitatively measure animal responses to BZs. We used a range of doses for each BZ compound to define response curves and found that all five of the alleles conferred resistance to BZ compounds equal to a loss of the entire beta-tubulin gene. These results prove that the parasite beta-tubulin alleles cause resistance. The E198V allele is found at low frequencies in natural parasite populations, suggesting that it could affect fitness. We performed competitive fitness assays and demonstrated that the E198V allele reduces animal health, supporting the hypothesis that this allele is less fit in field populations. Overall, we present a powerful platform to quantitatively assess anthelmintic resistance and effects of specific resistance alleles on organismal fitness in the presence or absence of the drug.

**Highlights:**

- All three canonical parasitic nematode beta-tubulin alleles (F167Y, E198A, F200Y) and two newly identified alleles (E198V, E198L) confer equal levels of benzimidazole resistance in a defined genetic background using single-generation, high-replication drug response assays.
- Beta-tubulin variants are strongly selected in albendazole conditions in multigenerational competitive fitness assays, but these alleles confer different levels of benzimidazole resistance over time.
- Only the E198V allele confers a fitness cost in control (non-benzimidazole) conditions as compared to all other tested beta-tubulin alleles, suggesting that this intermediate allele might only be found in field populations at low frequency because it causes reduced fitness.

**Graphical Abstract:** 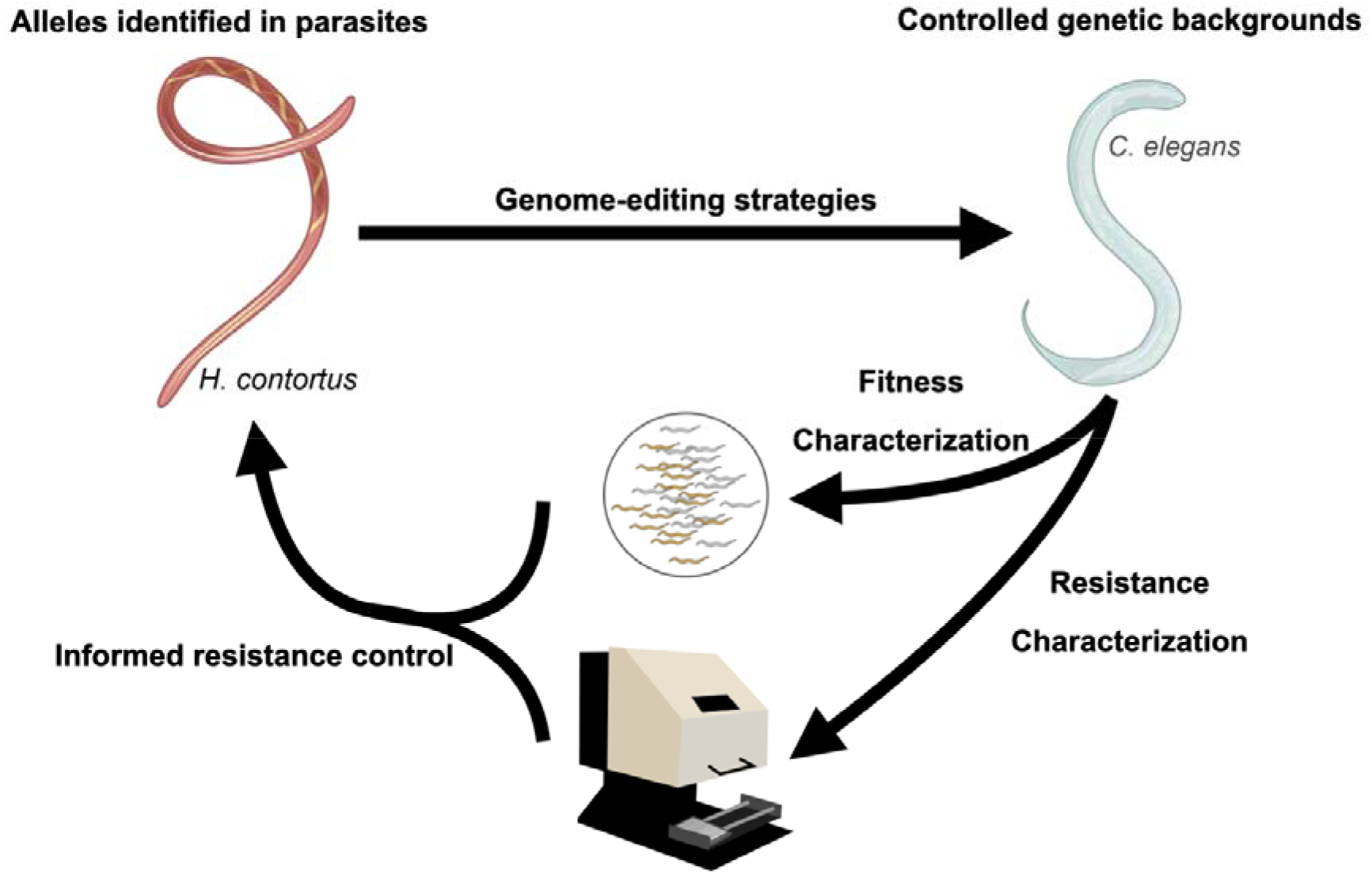

## 1. Introduction

Parasitic nematode infections cause a major health and economic burden that impacts billions of people around the globe (Hotez et al., 2014; Lustigman et al., 2012). In animal health, parasite infections lead to major reductions in livestock yields (Charlier et al., 2009; Kahn and Woodgate, 2012; Sutherland and Leathwick, 2011). Anthelmintics are the main defense against parasitic nematode infections, but an alarmingly low number of drug classes are both efficacious and safe (Kaplan and Vidyashankar, 2012). The four most commonly used classes of anthelmintics are the benzimidazoles (BZs), macrocyclic lactones (MLs), nicotinic acetylcholine receptor agonists (NAChA), and amino-acetonitrile derivatives (AADs). The first BZ was developed and implemented in veterinary medicine in 1962 under the name thiabendazole (Campbell and Cuckler, 1962; Merck and Sharp & Dohme Research Laboratories. Animal Science Research, 1962). Large-scale adoption and blanket usage of this drug placed a strong active pressure on parasites and quickly caused resistant populations throughout the world (Theodorides et al., 1970). Today, resistance to the two most widely used BZs, albendazole and fenbendazole, is common and found globally (Kaplan and Vidyashankar, 2012).

The genes that underlie resistance can be used to inform parasitic nematode control practices by monitoring the spread of resistance alleles in parasite populations. The alleles present in a population can predict drug efficacy and enable the maintenance of sensitive refugia populations (Hodgkinson et al., 2019; Muchiut et al., 2018). For BZ compounds, the first resistance gene, encoding a beta-tubulin, was identified in studies of the fungus *Aspergillus nidulans* (Hastie and Georgopoulos, 1971; Sheir-Neiss et al., 1978). Subsequently, *Caenorhabditis elegans* BZ selection experiments isolated resistant strains with mutations in a nematode-specific beta-tubulin gene, *ben-1 (Driscoll et al., 1989)*. This connection between beta-tubulin and nematode BZ resistance led to the discovery of mutations in *ben-1* homologs in BZ resistant parasitic nematode populations (Roos et al., 1990). The first allele in a *ben-1* homolog (*Hco-tub8-9*) that was correlated with BZ resistance in *Haemonchus contortus* populations was F200Y (Kwa et al., 1993, 1994). Subsequent surveys of resistant *H. contortus* populations identified two additional alleles correlated with resistance, F167Y (Silvestre and Cabaret, 2002) and E198A (Ghisi et al., 2007). These three putative resistance alleles are considered canonical because they are commonly found in resistant populations (Avramenko et al., 2019; Hinney et al., 2020; Redman et al., 2015). Recently, improved nematode genotyping techniques have enabled the detection of additional mutations at the 198 position: E198V, E198L, E198K, E198I, and E198Stop (Avramenko et al., 2019; Mohammedsalih et al., 2020; Redman et al., 2015; von Samson-Himmelstjerna et al., 2009). These beta-tubulin alleles represent promising markers of BZ resistance across parasitic nematode populations.

However, parasitic nematode studies have only correlated beta-tubulin alleles with BZ resistance. To go beyond correlation, a causal relationship between beta-tubulin and BZ resistance must be defined, which is difficult, if not impossible, in parasitic nematode species because genome-editing and characterized genetic backgrounds are required. Causal relationships between genes and phenotypes require empirical tests of sufficiency and necessity. To show that the beta-tubulin gene is sufficient for BZ response, functional replacement of the mutated gene must be shown to confer BZ sensitivity. This experiment was performed previously using overexpression of *C. elegans ben-1* or *H. contortus tub8-9 (Kwa et al., 1995)*. To show that beta-tubulin is necessary for BZ response, only the beta-tubulin gene must be mutated in a defined genetic background and shown to cause BZ resistance. Recently, the F167Y and F200Y alleles were shown to confer BZ resistance in *C. elegans* in a controlled genetic background (Hahnel et al., 2018; Kitchen et al., 2019).

Recent results indicate that BZ resistance is heterogeneous (Crook et al., 2016; Howell et al., 2008). Population-wide sequencing of beta-tubulin alleles have found different alleles in the same population (Avramenko et al., 2019; Hinney et al., 2020; Mohammedsalih et al., 2020; Redman et al., 2015; von Samson-Himmelstjerna et al., 2009). The frequency of each allele likely is determined both by its level of resistance and its effect on growth. Field observations have shown that populations maintain resistance after BZ pressure is removed (van Wyk et al., 1997), which suggests that they do not cause decreases in fitness. Measurements of the exact levels of resistance and any growth effects associated with the alleles are nearly impossible in field populations, so controlled laboratory experiments are required to test these hypotheses. In *C. elegans*, the F200Y and a loss-of-function *ben-1* allele conferred strong BZ resistance but no growth consequences compared to a wild-type control strain in conditions that lacked BZ (Hahnel et al., 2018), suggesting that the maintenance of resistance is not a detriment to fitness. One untested hypothesis of a possible interplay between resistance and fitness consequences was observed at the 198 position. The E198V allele has been detected at low frequencies in populations that also harbor the E198L allele (Avramenko et al., 2019; Mohammedsalih et al., 2020), suggesting that the E198V allele (**Supplemental Fig. 1**) confers lower levels of resistance and/or has a fitness detriment. In *C. elegans*, this interplay between resistance and fitness consequences can be determined using a defined genetic background, genome editing, and competitive fitness assays.

Here, we introduced beta-tubulin alleles from parasitic nematodes into a BZ-susceptible *C. elegans* laboratory strain using the CRISPR-Cas9 system. Independently generated alleles were assayed using multiple high-throughput techniques to determine levels of resistance and effects on fitness in both albendazole and fenbendazole. We definitively demonstrated that the beta-tubulin alleles from parasitic nematodes confer equivalent levels of BZ resistance. Additionally, we quantitatively assessed effects on competitive fitness in both control and BZ conditions. All parasite beta-tubulin alleles conferred BZ resistance as shown by an increase in allele frequency when competed against a wild-type control strain. In control conditions, each parasite beta-tubulin allele grew as well as a wild-type control strain, except for the fitness detriment conferred by the E198V allele. These results offer an explanation for why some alleles are found at higher frequencies and why the E198V allele is only found at low frequencies in parasite populations.

## 2. Methods

### 2.1 *C. elegans* strains

Animals in this study were grown at 20°C on modified nematode growth medium (NGMA) that contains 1% agar and 0.7% agarose seeded with OP50 bacteria (Andersen et al., 2014). For each assay, strains were grown for three generations following starvation in an attempt to reduce the multigenerational effects of starvation (Andersen et al., 2014). The existing strains harboring the *ben-1* deletion allele (*ean64*) or the F200Y alleles (*ean100*, *ean101*) (Hahnel et al., 2018) were used as controls for all experiments in this study. The barcoded wild-type control strain PTM229 *dpy-10(kah81)* was used in competitive fitness assays (Zhao et al., 2018). All strains in this study (Supplemental Table 1) were generated in the N2 genetic background with modifications introduced using the CRISPR-Cas9 genome-editing system as described previously (Hahnel et al., 2018).

### 2.2 CRISPR-Cas9 genome editing

All edited *ben-1* strains were generated in the N2 genetic background using CRISPR-Cas9-mediated genome editing. Targeted genome editing was facilitated using a co-CRISPR strategy (described below) to increase the chance of discovering successful edits (Kim et al., 2014). For the *ben-1* E198A, F167Y, E198V, and E198L allele replacement strains, we designed sgRNAs that targeted the *ben-1* locus and the *dpy-10* locus using the CRISPR design tool on the online analysis platform Benchling (www.benchling.com). These sgRNAs were ordered from Synthego (Redwood City, CA) and injected at 5 μM for the *ben-1* sgRNA and 1 μM for the *dpy-10* sgRNA. Single-stranded oligodeoxynucleotides (ssODN) templates for homology-directed repair (HDR) for both the *ben-1* and *dpy-10* alleles were ordered as ultramers (IDT, Skokie, IL) and injected at final concentrations of 6 μM and 0.5 μM, respectively. Cas9 protein (IDT, Skokie, IL) was also included and injected at a final concentration of 5 μM and incubated with the previously described sgRNAs and ssODNs at room temperature for one hour prior to injection. After incubation, the mixture was injected into the germlines of young adult hermaphrodite animals. These animals were singled to fresh 6 cm NGMA plates 18 hours post-injection. After 48 hours, F1 progeny were screened for animals with the Rol phenotype, and the individuals that displayed the Rol phenotype were singled onto new 6 cm NGMA plates and allowed to lay offspring. For the *ben-1* allele replacements, ssODN templates contained the desired nucleotide edit and a synonymous edit in the PAM site to prevent repeated cleavage by the sgRNA:Cas9 complex post introduction of the edit. In addition to preventing repeated cleavage at the PAM site, this edit introduced a new restriction site into the sequence. The E198A and E198V repair ssODNs introduced a *BtsC*I restriction site, the E198L repair ssODN introduced a *Hpy*188I restriction site, and F167Y introduced a *Sac*I cut site. Following PCR of the edited region, the PCR products were incubated with the restriction enzyme (New England Biolabs, Ipswich, MA) that had a recognition site introduced in the repair oligonucleotide for the desired edit. Successful edits were identified by an altered restriction pattern. F2 non-Rol individuals from parents with successful edits were placed on single plates to generate homozygous progeny. F2 individuals were then genotyped following the same process described above, and successful edits were Sanger sequenced to ensure homozygosity of the desired edits before use in future experiments. Independent edits of each allele were generated to control for off-target effects of the CRISPR-Cas9 process. All oligonucleotides used in this study are available upon request (Supplemental Table 2).

### 2.3 High-throughput fitness assays

High-throughput fitness assays were performed as previously described (Evans and Andersen, 2020; Zdraljevic et al., 2017). Briefly, a 0.5 cm^3^ NGMA piece was cut from a plate containing starved individuals to a new NGMA plate. Two days later, gravid hermaphrodites from the new NGMA plate were placed into a bleach solution on a 6 cm agar plate to clean and synchronize each strain. The next day, five L1 larvae were moved to 6 cm NGM plates and allowed to reproduce over the course of five days until the offspring had reached the L4 larval stage. Five L4 individuals were transferred to new 6 cm NGMA plates and allowed to grow and reproduce for four days. After this growth, strains were bleach synchronized to generate a large pool of synchronized unhatched embryos for each strain. These embryos were diluted to roughly one embryo per μL in K medium (Boyd et al., 2012). Embryos were placed into 96-well plates at roughly 50 embryos per well in 50 μL of K medium and allowed to hatch overnight. The next morning, a mixture HB101 lysate (García-González et al., 2017) at a concentration of 5 mg/mL and either drug (albendazole or fenbendazole) in DMSO or DMSO alone was added to the solution. Final DMSO concentrations were kept at 1% for all doses and experiments. For dose response assays, drugs were used at 0, 6.25, 12.5, 25, 50, and 100 μM. For high-replication albendazole and fenbendazole assays, both drugs were used at a final concentration of 30 μM because at this concentration N2 showed a severe developmental delay and the parasite beta-tubulin alleles had begun to be developmentally delayed in dose response experiments. After 48 hours of growth in either control or BZ conditions, animals were scored using the COPAS BIOSORT (Union Biometrica, Holliston MA) after treatment with sodium azide at 50 mM to straighten the animals (Brady et al., 2019; Hahnel et al., 2018; Zdraljevic et al., 2017). Animal optical density integrated by the animal length (EXT) was measured with the COPAS BIOSORT for each individual. The median optical density (median.EXT) of nematode populations in each well was used for quantitative BZ responses.

### 2.4 Optical density trait processing

All growth trait processing and analyses were completed using the R(3.6.1) package *easysorter (Shimko and Andersen, 2014)* with the v3_assay = TRUE argument added to the *sumplate* function. Beyond modifications for changes in the assay workflow, analyses were performed as described previously (Hahnel et al., 2018) for high-replication single-concentration experiments. For dose-response experiments, phenotypes for each strain were normalized to the value of each strain in control conditions (1% DMSO). For normalization, we calculated the mean of our trait of interest (median.EXT) in control conditions and subtracted that value from the phenotypic values from each of the different doses. This normalization was performed on a strain-specific basis to control for any differences in the starting populations among strains.

### 2.5 Competition assays

We used previously established pairwise competition assays to assess organismal fitness (Zhao et al., 2018). The fitness of a strain is determined by comparing the allele frequency of a strain against the allele frequency of a wild-type control strain. Both strains harbor molecular barcodes to distinguish between the two strains using oligonucleotide probes complementary to each barcode allele. In short, ten L4 individuals of each strain were placed onto a single 6 cm NGMA plate along with ten L4 individuals of the PTM229 strain (an N2 strain that contains a synonymous change in the *dpy-10* locus that does not have any growth effects compared to the normal laboratory N2 strain (Zhao et al., 2018)). Ten independent NGMA plates of each competition were prepared for each strain in each test condition, control (DMSO) or BZ (albendazole and DMSO). We prepared ten replicates of each competition, except the deletion allele of *ben-1,* which only had six replicates of the control condition because of contamination on four of the plates. The N2, F200Y (*ean101*), and *ben-1* deletion (*ean64*) strains were included to ensure that assays were reproducible and that all plates had an effective albendazole concentration (Hahnel et al., 2018). Plates were grown for roughly one week to starvation. Animals were transferred to a new plate of the same condition by transferring a 0.5 cm^3^ NGMA piece from the starved plate onto the new plate. The remaining individuals on the starved plate were washed into a 15 mL Falcon tube with M9 buffer, concentrated by allowing the animals to settle for roughly one hour, and stored at −80°C. DNA was extracted using the DNeasy Blood & Tissue kit (Qiagen 69506). Competitions were performed for seven generations, and animals were collected after generations, one, three, five, and seven.

We quantified the relative allele frequency of each strain as previously described (Zhao et al., 2018). In short, a digital PCR approach with TaqMan probes (Applied Biosciences) was used. Extracted genomic DNA was digested with *Eco*RI for 30 minutes at 30°C, purified with Zymo DNA cleanup kit (D4064), and diluted to approximately 1 ng/μL. Using TaqMan probes as described previously (Zhao et al., 2018), the digital PCR assay was performed with a Bio-Rad QX200 device with standard probe absolute quantification settings. The TaqMan probes bind to wild-type *dpy-10* and the *dpy-10* allele present in PTM229 selectively. Relative allele frequencies of each tested allele were calculated using the QuantaSoft software and default settings. Calculations of relative fitness were calculated by linear regression analysis to fit the data to a one-locus generic selection model (Zhao et al., 2018).

### 2.6 Statistical analyses

All statistical comparisons were performed in *R* version 3.6.1 (Core Team and Others, 2013) with the *Rstatix* package. The *tukeyHSD* function was used on an ANOVA model with the formula *phenotype ~ strain* to calculate differences between population trait values. The EC50 values for the fenbendazole dose response assay were calculated using the *drm* function from the *drc* package with fct = LL.3 (Ritz et al., 2015).

### 2.7 Research Data

S1 contains dose-response curve data for albendazole treatment. S2 contains regressed data at high-replication for albendazole treatment for canonical alleles. S3 contains regressed data at high-replication for albendazole treatment for newly identified alleles. S4 contains dose-response curve data for fenbendazole treatment. S5 contains regressed data at high-replication for fenbendazole treatment for canonical alleles. S6 contains regressed data at high-replication for albendazole treatment for newly identified alleles. S7 contains regressed data at high-replication for fenbendazole treatment for all alleles. S8 contains raw phenotype data for DMSO conditions. S9 contains calculated EC50 values for each allele. S10 contains the calculated allele frequency for the competitive fitness assay in albendazole and DMSO. S11 contains calculated competitive fitness values in albendazole and DMSO. S12 contains allele designation to ECA translations. S13 is an xlsx file with raw ddPCR data from the competitive fitness assay. Supplemental Table 1 contains a list of all strains along with genotypes and strain designations. Supplemental Table 2 contains all oligonucleotides used in the study for CRISPR, PCR, and allele frequency calculations. All tables, datasets, along with code for analysis and generation of figures can be found at GitHub (https://github.com/AndersenLab/ben1_2020_CMD).

## 3. Results

### 3.1 All three canonical parasitic nematode beta-tubulin alleles confer BZ resistance

We performed BZ dose-response assays on strains with the three canonical parasitic nematode beta-tubulin alleles (F200Y, F167Y, and E198A) along with positive and negative controls for albendazole effects (N2 and the *ben-1* deletion allele, respectively). All strains were generated in the N2 background, so the allele at the *ben-1* locus is the only difference between the genome-edited strain and the unedited parent strain. The BZ response assays used a flow-based device to measure the optical densities of hundreds of animals in each of 12 replicate cultures per strain (see Methods). Optical density refers to the amount of the laser light that is obstructed by an individual integrated over the amount of time that the individual interrupts the laser. Optical density was used to quantify BZ responses as a proxy for nematode health because *C. elegans* grows longer and more dense as it develops. A decrease in optical density represents a developmental delay, which is commonly observed in responses to BZ treatments (Driscoll et al., 1989; Hahnel et al., 2018; Zamanian et al., 2018). This experimental setup allowed us to determine whether the parasitic nematode beta-tubulin alleles quantitatively caused resistance as compared to the sensitive wild-type *ben-1* allele and the resistant *ben-1* deletion allele (**Fig. 1**). We tested the BZ responses of these strains to either albendazole or fenbendazole. We also tested for developmental defects in control conditions, which would indicate detrimental fitness effects, but did not detect any defects. Surprisingly, the F167Y strain developed faster than the wild-type strain in control conditions (**Supplemental Fig. 2**).

**Fig. 1.**
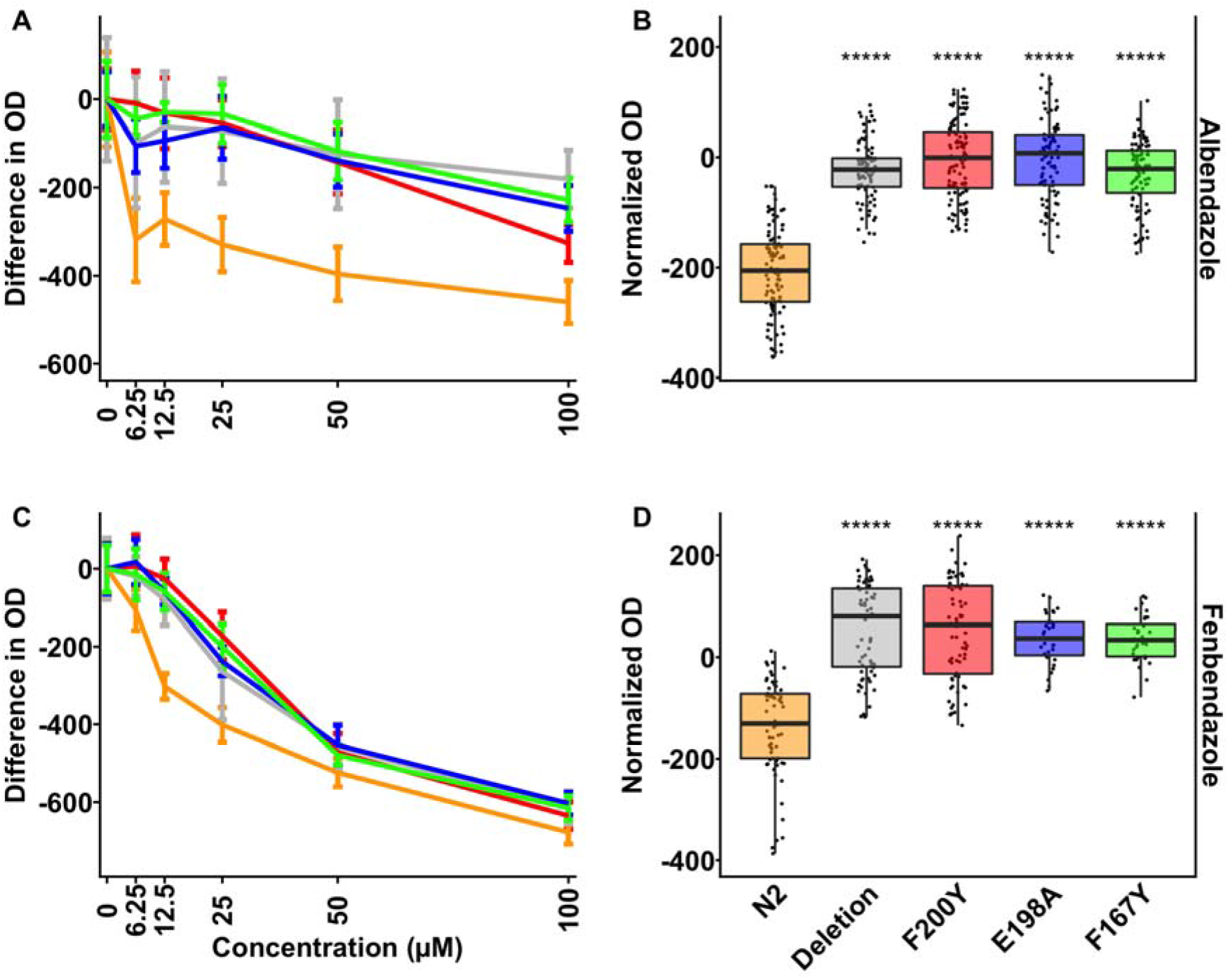
Quantitative responses of canonical parasite beta-tubulin alleles to albendazole and fenbendazole. Drug-response assays for three canonical parasite beta-tubulin alleles, F200Y, E198A, and F167Y, are shown. (A, C) Normalized values were calculated by subtracting the mean optical density in a strain from each population measurement. Normalized trait values are shown in response to six drug concentrations (0, 6.25, 12.5, 25, 50, and 100 μM) of BZ. (B, D) Regressed optical density values of responses to 30 μM BZ at high-replication are shown on the y-axis. Each point represents a regressed optical density value calculated per well containing approximately 35-50 animals. Significant differences between N2 and all other strains are shown as asterisks above the data from each strain (p < 0.001 = ***, p <0.0001 = ****, Tukey HSD). (A) and (B) show responses to albendazole; (C) and (D) show responses to fenbendazole.

Albendazole and fenbendazole caused different phenotypic effects on the set of *C. elegans* strains. In response to albendazole (**Fig. 1A,B**), the wild-type laboratory strain was sensitive, measured as a large decrease in development rate (decreased animal optical density) even at the lowest concentration of albendazole. As concentrations increased, the laboratory strain was only marginally more affected. By contrast, the *ben-1* deletion strain was resistant at all concentrations, including the highest concentrations of albendazole. The three canonical parasitic nematode beta-tubulin alleles had dose-response curves statistically similar to the *ben-1* deletion strain, which suggests they conferred equal levels of resistance. To determine subtle albendazole effects among the strains, we measured development in the presence of 30 μM albendazole for each strain at high replication. The three canonical parasitic nematode beta-tubulin strains were statistically indistinguishable from the *ben-1* deletion strain and were significantly more resistant than the wild-type strain (**Fig. 1B**).

Fenbendazole caused more drastic changes to *C*. *elegans* development in a dose-dependent manner (**Fig. 1C,D**). The wild-type laboratory strain showed significant developmental delays across the range of doses studied. Unlike albendazole, each of the *ben-1* mutant alleles (including the *ben-1* loss-of-function deletion allele) also showed developmental defects as the concentration of fenbendazole increased. At the 100 μM fenbendazole dose, all of the strains were severely developmentally delayed. To determine subtle fenbendazole effects among the strains, we measured growth in the presence of 30 μM fenbendazole for each strain at high replication. Again, the three canonical parasitic nematode beta-tubulin strains responded similarly to the *ben-1* deletion strain and were less sensitive to fenbendazole as compared to the wild-type control strain (**Fig. 1D**). For each genome-edited allele, we tested two independently generated strains, and all independent strain duplicates showed similar responses across all doses of both BZ compounds and at high-replication (**Supplemental Fig. 3, Supplemental Fig. 4**).

The results of the dose-response and the high-replication assays indicated that the canonical parasitic nematode beta-tubulin alleles conferred resistance to BZs (**Fig. 1, Supplemental Fig. 5**). The wild-type strain had a lower EC50 than any of the canonical parasitic nematode beta-tubulin alleles, because it is more sensitive to BZ treatment. The EC50 values for the canonical parasitic nematode beta-tubulin alleles were statistically the same, which suggested that these alleles conferred the same quantitative level of resistance. We could not calculate EC50 values in albendazole because the full range of phenotypic responses required higher concentrations than tested. Regardless, the change in responses among the canonical parasitic nematode beta-tubulin alleles and the susceptible wild-type strain was apparent. These results demonstrated that each of the three canonical parasitic nematode beta-tubulin alleles caused resistance to albendazole and fenbendazole.

### 3.2 Two of the less frequently observed parasitic nematode beta-tubulin alleles also confer BZ resistance

Other beta-tubulin alleles beyond the canonical alleles have been identified using population sequencing approaches (Avramenko et al., 2019; Mohammedsalih et al., 2020; Redman et al., 2015). These alleles are not present in all parasites in a population, so they might confer different levels of BZ resistance and/or effects on growth. We introduced the E198V and E198L alleles into the N2 genetic background to perform quantitative drug response assays using the same flow-based measurement and experimental design as previously described. In response to albendazole, both the E198V and E198L strains responded similarly to the *ben-1* deletion strain at each BZ concentration (**Fig. 2A**). Next, we performed a high-replication experiment to determine if the edited beta-tubulin alleles conferred small differences in albendazole resistance at 30 μM. As previously observed with the canonical parasitic nematode beta-tubulin alleles, both E198V and E198L showed statistically the same response to albendazole as the *ben-1* deletion strain (**Fig. 2B**). In fenbendazole, the results followed the same pattern as observed in the canonical parasitic nematode beta-tubulin alleles (**Fig. 2C,D**). Except for the highest fenbendazole concentrations, both E198V and E198L were developmentally delayed but to a lesser extent than the susceptible N2 strain (**Fig. 2C**). At 50 μM fenbendazole, N2 and the mutant *ben-1* strains were statistically indistinguishable from each other and all showed significant developmental delays. At high-replication, E198V and E198L were statistically more resistant than N2 (p < 0.0001, Tukey HSD) but not different from the *ben-1* deletion strain at 30 μM fenbendazole (**Fig. 2D**). All of the tested beta-tubulin alleles conferred the same fenbendazole response as shown by the EC50 estimates (**Supplemental Fig. 5**). Similar to what we found with the canonical parasitic nematode beta-tubulin alleles, the two independently generated genome-edited strains for each of the E198V and E198L alleles were not significantly different from each other in the dose-response assay (**Supplemental Fig. 3**) or the high-replication single-dose assay (**Supplemental Fig. 4**). Additionally, E198V and E198L were tested in control conditions and showed no significant differences from the N2 strain, which suggested that they do not cause viability defects in this single-generation assay (**Supplemental Fig. 2**).

**Fig. 2.**
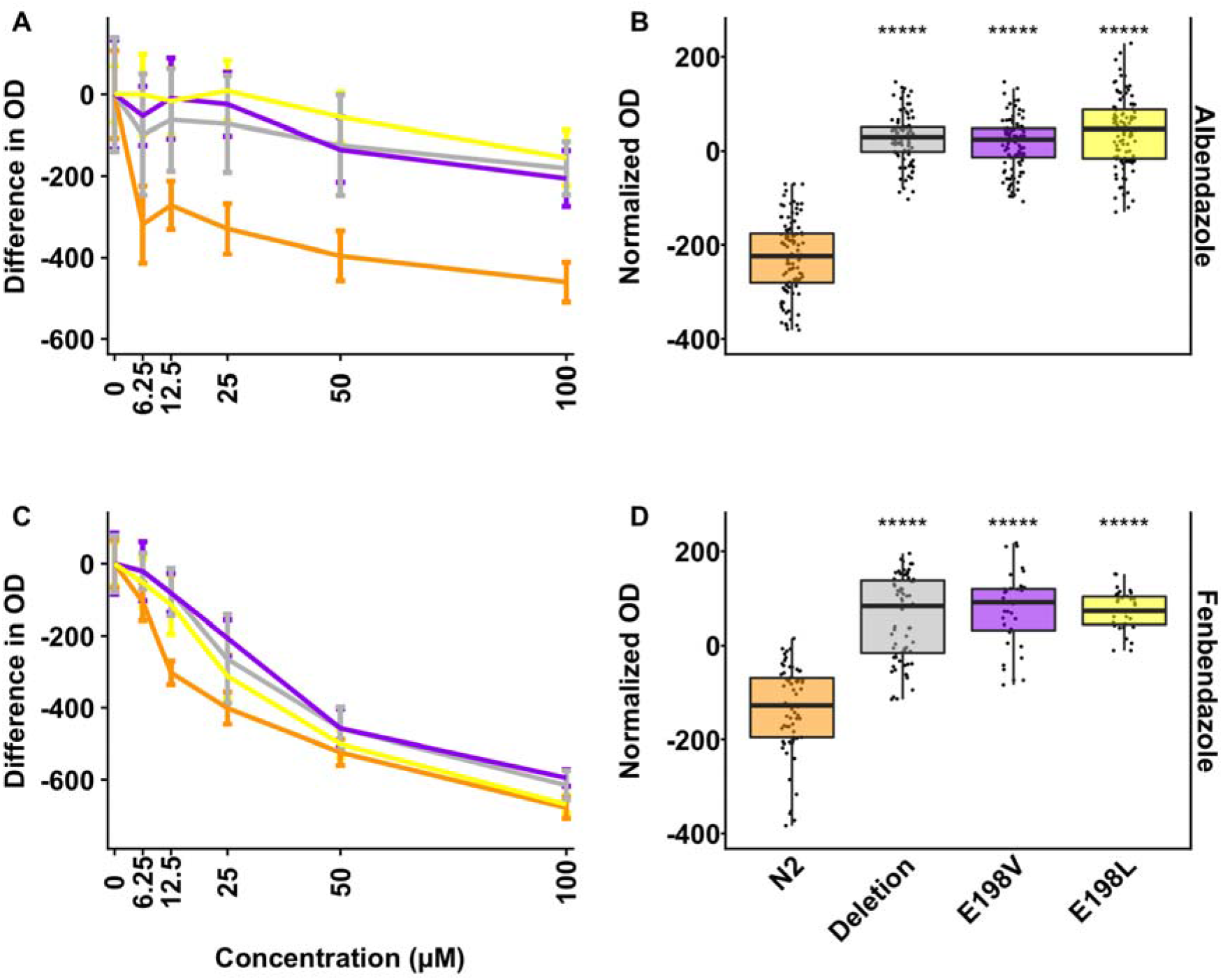
Quantitative responses to albendazole and fenbendazole conferred by the E198V and E198L alleles. Drug-response assays for two of the non-canonical parasitic nematode beta-tubulin alleles, E198V and E198L, are shown. (A, C) Normalized values were calculated by subtracting the mean trait value measured for a population of individuals in the control condition (0 μM BZ) from the trait values measured for a population of individuals at all BZ concentrations (0, 6.25, 12.5, 25, 50, and 100 μM). (B, D) Regressed optical density values of responses to 30 μM BZ at high-replication are shown on the y-axis. Each point represents a regressed optical density value calculated per well containing approximately 35-50 animals. Significant differences between N2 and all other strains are shown as asterisks above the data from each strain (p < 0.001 = ***, p <0.0001 = ****, Tukey HSD). (A) and (B) show responses to albendazole; (C) and (D) show responses to fenbendazole.

### 3.3 Multi-generational competitive fitness assays quantify albendazole selection and growth effects caused by beta-tubulin alleles

We performed multi-generational assays to measure competitive fitness effects between strains with the parasitic nematode beta-tubulin alleles and the wild-type *ben-1* allele. This assay is more sensitive than our previous single-generation high-throughput assays because differences in reproductive rate also influence fitness. Additionally, small changes in fitness are more easily observed as they accumulate over multiple generations. If an allele confers a deleterious fitness effect compared to the wild-type strain, then that strain will decrease in frequency over the generations of the competition assay. Conversely, if an allele confers a beneficial effect compared to the wild-type allele, then that strain will increase in frequency over the generations of the competition assay. Finally, if an allele has no difference in effect compared to the wild-type allele, then the two strains would be found at approximately equal frequencies over the generations of the competition assay. The N2, *ben-1* deletion, and the F200Y strains have been tested in both albendazole and control conditions previously (Hahnel et al., 2018). The wild-type strain showed no differences in competitive fitness in either control or albendazole conditions compared to the barcoded wild-type control strain (**Fig 3**). As demonstrated previously (Hahnel et al., 2018), the F200Y and the *ben-1* deletion strains had no fitness consequences in control conditions but had growth advantages in BZ conditions. In albendazole conditions, we expected that all parasite beta-tubulin alleles would confer resistance in multi-generational assays because we observed resistance in the high-throughput assays (**Fig. 1, Fig. 2**). Therefore, the frequencies of the parasite beta-tubulin alleles should increase over time in albendazole conditions. As expected, all edited alleles showed a steady increase in frequency (as compared to the wild-type strain) across each subsequent generation in albendazole conditions (**Fig. 3A**). The allele frequencies as compared to the wild-type control strain can be used to calculate competitive fitness. We found that each edited allele conferred significant competitive advantages in albendazole conditions (**Fig. 3B**). However, the E198V allele was significantly more developmentally delayed than the F200Y allele across the seven generations of albendazole selection (p < 0.01, Tukey HSD) (**Fig. 3A,B**).

**Fig. 3.**
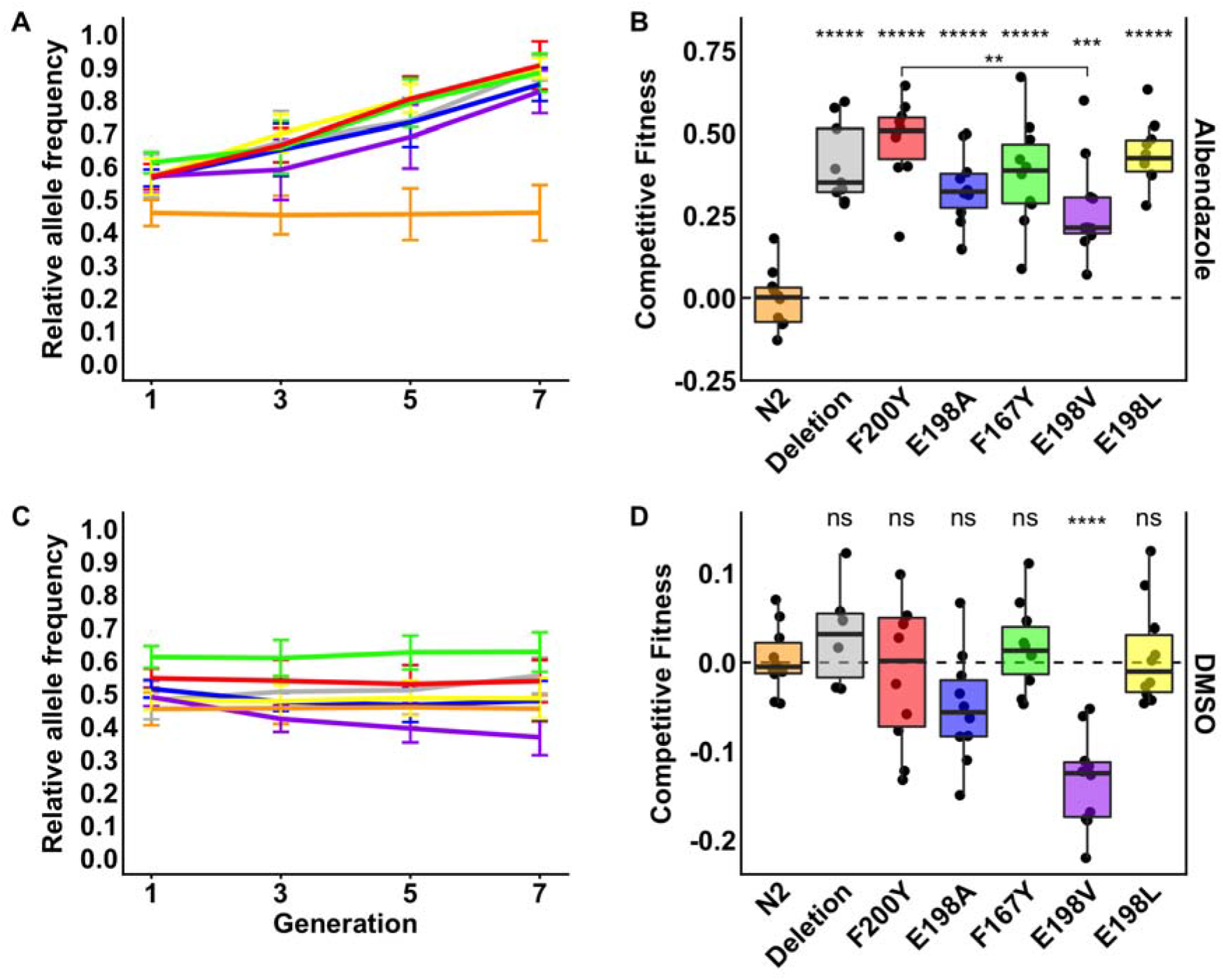
Variation in competitive fitness among *ben-1* alleles in both albendazole and control conditions. (A) Results from a multi-generational competition assay between a barcoded N2 strain and a strain of each edited beta-tubulin strain in albendazole conditions. Generation number is shown on the x-axis, and relative allele frequencies are plotted on the y-axis. (B) Tukey box plots of log_2_-transformed competitive fitness values in albendazole conditions are shown. The tested allele is shown on the x-axis, and competitive fitness values as compared to the wild-type strain are plotted on the y-axis. Each point represents the competitive fitness of a biological replicate. Significant differences between N2 and all other strains are shown as asterisks above the data from each strain (p < 0.001 = ***, p <0.0001 = ****, Tukey HSD). The level of significance between the F200Y and E198V strains is shown as a bracket (p <0.01, Tukey HSD). (C) Results from multi-generational pairwise competition assays in DMSO control conditions are plotted as in (A). (D) Tukey box plots of competitive fitness in control DMSO conditions are shown as in (B). Significant differences between N2 and all other strains are shown as asterisks above the data from each strain (p <0.0001 = ****, Tukey HSD).

Because the multi-generational assays might be more sensitive than the high-throughput assays, we wanted to compete strains in control conditions to investigate whether any parasite beta-tubulin alleles confer fitness disadvantages without the presence of drug selection. We found no deleterious effects in the high-throughput assays, so we expected that all alleles would be equally fit in the multi-generational assays. In control conditions, only the E198V allele conferred a difference in growth, decreasing in abundance compared to the wild-type strain over seven generations (**Fig. 3C**). Based on the competitive fitness values, this strain was significantly less fit than the wild-type strain in control growth conditions (p<0.0001, Tukey HSD) (**Fig. 3D**). This decrease suggests that strains with the E198V allele can be outcompeted in normal conditions by strains with the wild-type allele at the *ben-1* locus.

## 4. Discussion

### 4.1 Parasitic nematode beta-tubulin alleles confer BZ resistance

After exposure to BZ compounds, the frequencies of certain beta-tubulin alleles increase in parasitic nematode populations, including F167Y, E198A, E198I, E198K, E198L, E198V, E198Stop, and F200Y (Avramenko et al., 2019; Ghisi et al., 2007; Kwa et al., 1994; Mohammedsalih et al., 2020; Redman et al., 2015; von Samson-Himmelstjerna et al., 2009; Silvestre and Cabaret, 2002). These alleles have long been hypothesized to cause BZ resistance, but the lack of controlled genetic backgrounds and genome-editing tools have made it difficult to test this hypothesis in parasitic nematode species. Previous studies have used *C. elegans* overexpression experiments to show that beta-tubulin is sufficient to restore BZ susceptibility (Kwa et al., 1995). Since the introduction of the CRISPR-Cas9 system in *C. elegans*, experiments on gene function at the single amino acid level have become possible. We used genome editing to independently introduce five parasitic nematode beta-tubulin alleles, F167Y, E198A, E198L, E198V, and F200Y, into the *C. elegans ben-1* locus in a defined genetic background. We tested each of these alleles using both high-replication growth assays and competitive fitness assays in BZ and control conditions. We found all alleles conferred the same level of resistance to albendazole and fenbendazole in the high-replication growth assays that encompass a single generation. However, we found in multi-generational competitive fitness assays that the E198V allele confers lower resistance in albendazole conditions than the F200Y allele. Additionally, the E198V allele also had decreased fitness compared to the wild-type strain in control conditions. Our results definitively prove that these beta-tubulin alleles confer BZ resistance and that some of these alleles can also affect fitness in laboratory conditions.

The two assays used in this study have different advantages and disadvantages. The high-replication high-throughput assay provides an excellent platform for quickly quantifying if an allele confers resistance to BZs. The assay enables high levels of replication. One disadvantage of this assay is that traits are measured after only one generation of growth under drug selection, which could fail to identify subtle differences in fitness or resistance. By contrast, the multigenerational competition assay enables the measurement of small differences between the edited alleles and the wild-type strain over seven generations, which can detect subtle differences in fitness or resistance. Each generation compounds the fitness impacts of the edited alleles. These assays in combination provide a powerful platform to determine levels of resistance and potential fitness consequences.

Our ability to test beta-tubulin benzimidazole resistance alleles in a high-throughput and sensitive manner enables the study of additional alleles. In this study, we tested five of the eight alleles identified in parasite populations. The E198I, E198K, and E198Stop alleles still need to be tested in defined genetic backgrounds. In addition to testing if these alleles confer resistance, we can use the multigenerational competition assays to assess whether they confer any fitness effects. The results of these experiments can be used to implement proper management of resistance to benzimidazoles in field populations. For example, knowledge about quantitative levels of BZ resistance can alter anthelmintic treatment strategies if high BZ resistance alleles are detected in populations. Importantly, the field requires cheap, point-of-care diagnostic tests for validated resistance alleles. If we had a complete catalog of resistance alleles, veterinarians and farmers could more easily maintain refugia populations and control benzimidazole resistance.

### 4.2 Differential responses to albendazole and fenbendazole in *C. elegans*

Albendazole and fenbendazole are both widely used anthelmintic compounds that often treat similar parasitic nematode species. However, these compounds have different levels of effectiveness and safety on host species. For example, albendazole has previously been shown to have teratogenic effects in pregnant rats and sheep (Capece et al., 2003; Navarro et al., 1998). By contrast, the primary fenbendazole metabolite in ruminants, oxibendazole, showed no toxic effects on pregnant mice, rats, sheep, or cattle (Theodorides et al., 1977). In addition to differential responses in the host species treated with albendazole or fenbendazole, parasitic nematodes respond differently to these two compounds. A recent study found that, in response to albendazole, fewer genes are differentially expressed compared to fenbendazole (Stasiuk et al., 2019). Our study also found differential responses to albendazole and fenbendazole, where *C. elegans* was overall less susceptible to albendazole than fenbendazole. Only the highest albendazole concentrations caused detrimental effects on the strains with parasitic nematode beta-tubulin alleles. By contrast, fenbendazole caused detrimental effects on all strains at all concentrations. The difference in these responses suggests that future studies of beta-tubulin copy number, tissue-specificity, and specific drug interactions could be fruitful.

One possibility underlying this difference in response is that fenbendazole might have additional targets beyond beta-tubulin, including genes that encode beta-tubulin interacting proteins. Additionally, the *C. elegans* laboratory wild-type strain N2 has five beta-tubulin genes beyond *ben-1*, which could provide targets for fenbendazole. The two most highly expressed beta-tubulins, *tbb-1* and *tbb-2*, harbor the resistant tyrosine at the 200 position making them less than ideal candidates. However, it is possible that fenbendazole binds these beta-tubulins with lower affinity. Previous studies have discovered additional loci that underlie differential responses to fenbendazole (Zamanian et al., 2018), and the loci are not shared between albendazole and fenbendazole. These loci could be BZ compound specific. Moreover, studies of gene expression in response to different BZ compounds have shown that fenbendazole exposure causes more differentially expressed genes than albendazole (Stasiuk et al., 2019). Many of these genes encode enzymes essential for biotransformation, including cytochrome P450 family members, nuclear hormone receptors, and UDP-glucuronosyltransferases. To make more BZ compounds effective across more parasitic nematode species, we must better understand these BZ-specific effects.

### 4.3 Benzimidazole resistance supersedes detrimental fitness costs caused by beta-tubulin alleles

In the field, parasitic nematodes must balance responses to abiotic stresses, like anthelmintic drugs, and growth. For example, alleles that confer extremely high levels of resistance, but also cause severe detrimental fitness consequences, will be strongly selected when BZs are used on the population. However, these same alleles will quickly be selected against when the drug is removed. The selective pressure caused by the anthelmintic drug must be greater than the reduced fitness caused by the resistance allele when the drug is removed. Alleles that confer resistance and have no fitness consequences will be maintained in field populations. This argument could be one reason why we only find specific beta-tubulin alleles across parasite populations.

The multi-generational competitive fitness assays can be used to investigate this balance by comparing relative fitness in drug and control conditions. We found that all parasitic nematode beta-tubulin alleles conferred resistance to albendazole over seven generations when competed against wild-type animals (**Fig. 3**). However, some alleles caused different levels of resistance to albendazole. The E198V allele conferred significantly lower levels of albendazole resistance than the F200Y allele, and the E198V allele was also the only allele to confer detrimental fitness in control conditions. This results suggests that this allele could be less advantageous than other beta-tubulin alleles in that it causes less BZ resistance and decreased fitness. Matching these results, this allele is found at low frequencies in field populations (Avramenko et al., 2019). The E198V allele might be selected when BZ drugs are used, but when the drug is removed from a population, this allele would cause a fitness detriment and not be maintained in the population. Conversely, all of the other tested alleles showed no detrimental fitness consequences when the drug is not present. These results suggest that, in field populations with persistent BZ resistance after removal of treatment (van Wyk et al., 1997), the beta-tubulin allele might also persist because of the lack of fitness cost.

Our results can be explained by this balance between BZ resistance and fitness costs, but it is still not clear why parasite populations harbor low frequency E198V alleles and higher frequencies of the E198L allele (Avramenko et al., 2019; Mohammedsalih et al., 2020). The E198L allele can only be generated after changing two nucleotides, and the E198V allele is one of those changes (**Supplemental Figure 1**). Why would a parasite population make the two-step transition from wild-type to E198V to E198L, instead of a single step to one of the canonical alleles, like E198A? As we showed, the E198A allele is equally resistant to albendazole as E198L and neither alleles seems to confer a fitness cost (**Fig 3**). One explanation is that some or all beta-tubulin alleles confer dominant resistance to BZ compounds. It is not known the allelic status in field populations, and we only tested homozygous *C. elegans* strains in our assays. Heterozygous animals harboring the E198V allele might be resistant to BZ compounds, and other alleles might only be resistant when homozygous. In that way, resistance conferred in heterozygous individuals can be selected for additional changes that will change the E198V allele to an allele that does not cause a fitness cost. Dominance relationships for these alleles must be tested in characterized genetic backgrounds and sensitive BZ assays. Resistance conferred by each allele in a heterozygote could also vary, and the ordering of the resistance in heterozygotes could help us understand the speed at which resistance can spread through a population. Another explanation for this disparity in the frequencies of alleles across resistant populations could be the codon preference in the target species. Previous work found that codon preference can vary dramatically among species (Mitreva et al., 2006). If the codon preference of a species favors a transition to E198V instead of E198A that could help explain this transition. Another explanation could be suggested by a non-significant trend observed in our competitive fitness assay where we found that the E198A allele appeared to confer a fitness detriment as compared to F167Y, E198L, and F200Y in albendazole (**Fig 3A,B**). The E198A allele appears to more closely resemble the E198V allele in competitive fitness over seven generations than the other tested alleles. However, we require increased levels of replication to identify if this trend is significant.

The balance between BZ resistance and fitness costs are also influenced by the complement of beta-tubulin genes present in a species and the tissues and times in which they are expressed. For example, *C. elegans* has six beta-tubulin genes and some of these have been shown to be at least partially functionally redundant with each other (Ellis et al., 2004).Importantly, the two most highly expressed beta-tubulin genes are *tbb-1* and *tbb-2*, and both of these genes encode beta-tubulins that should be resistant to BZ treatments. *H. contortus* only has four beta-tubulin genes and the two most highly expressed are sensitive to BZ treatments (Saunders et al., 2013). This lack of redundancy and prevalent drug sensitivity could explain both the kinetics of resistance and the persistence of alleles across populations. Competitive fitness assays would be difficult to perform in *H. contortus* because of the previously discussed issues of lack of a controlled genetic background and genome-editing tools. However, one possibility to get around this obstacle is to make the *C. elegans* complement of beta-tubulin more like *H. contortus*. Removal of redundant beta-tubulin genes in *C. elegans* would study the effects of these alleles in a model closer to the situation in *H. contortus* populations. Overall, the balance parasites must maintain between anthelmintic resistance and maintenance of competitive fitness when drugs are not present is an extremely relevant approach to understand anthelmintic effectiveness.

### 4.4 Future avenues and methods of research

The approach described here shows that hypotheses generated from the study of parasitic nematode populations can be tested and measured using the model *C. elegans*. Future studies of homozygous and heterozygous resistance could impact parasitic nematode control in field populations. Additionally, the generation of more *H. contortus*- like models using *C. elegans* will allow for more accurate replication of the natural situations in which alleles exist. The continued interplay of research in *C. elegans* and parasite species allows for quick validation of resistance alleles and measurement of any fitness consequences associated with the identified alleles.

## Acknowledgements

We would like to thank Katie Evans, Robyn Tanny, Sam Widmayer, and Janneke Wit for their feedback and comments on this manuscript. C.M.D. was supported by the Biotechnology Training Program at Northwestern University (T32 GM008449). S.R.H. was funded by a DFG fellowship (HA 8449/1-1) from the Deutsche Forschungsgemeinschaft. E.C.A. was funded by the National Institutes of Health NIAID grant R21AI121836. This study would not have been possible without data from Wormbase, the *Caenorhabditis* Genetics Center (P40 OD010440), and the *Caenorhabditis elegans* Natural Diversity Resource (NSF CSBR 1930382). The funding sources had no impact on the design of this study. We would also like to thank Samantha Peters for her illustrations of *C. elegans* and *H. contortus*.

## Supplemental Figures

**Supplemental Figure 1.**
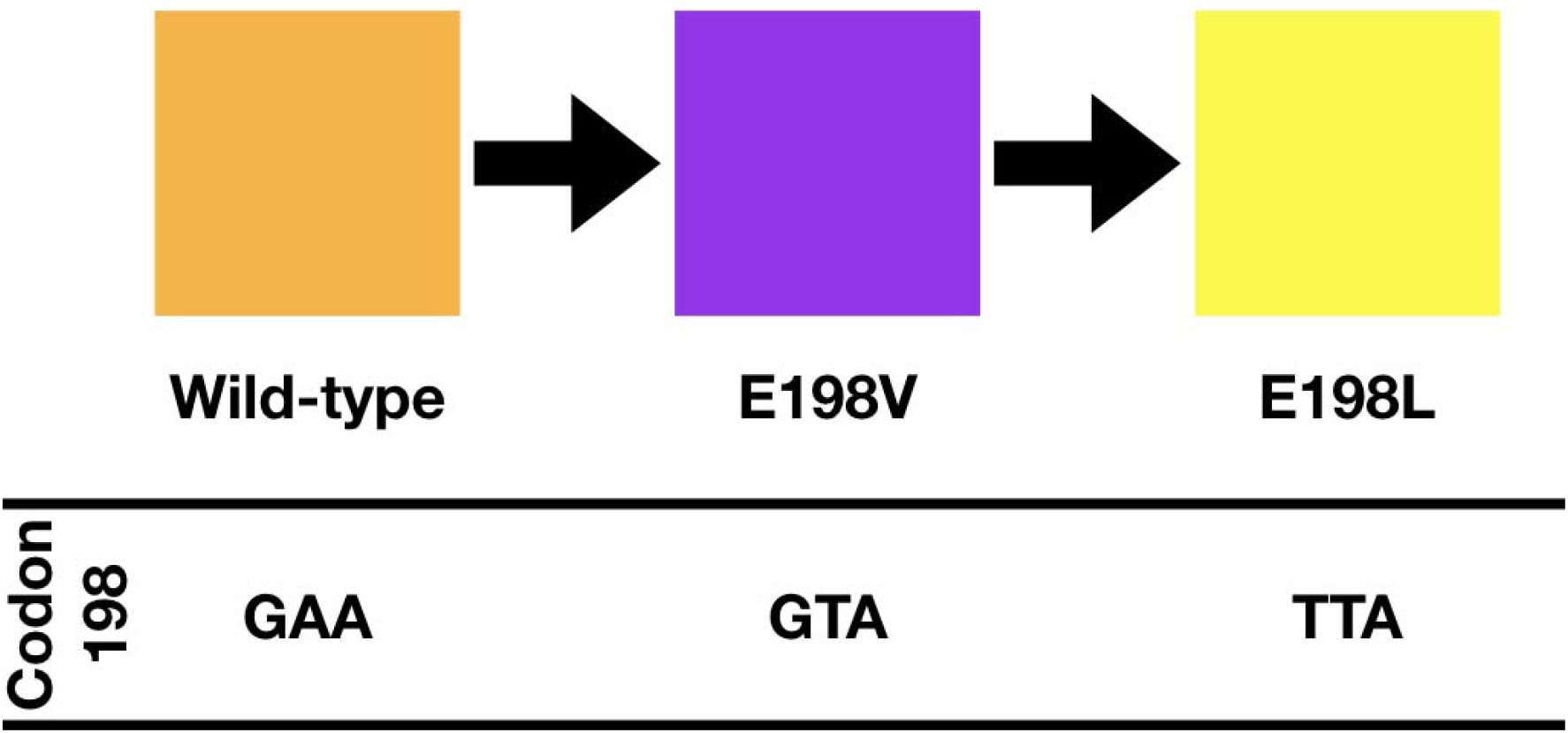
Graphical interpretation of the predicted resistance based on allele frequency observations in parasite populations (Avramenko et al., 2019). The boxes at the top represent the strains that are used in this study. The codon at position 198 is shown below and demonstrates how E198V (purple) could be an intermediate one nucleotide step from E198 (orange) to E198L (yellow).

**Supplemental Figure 2.**
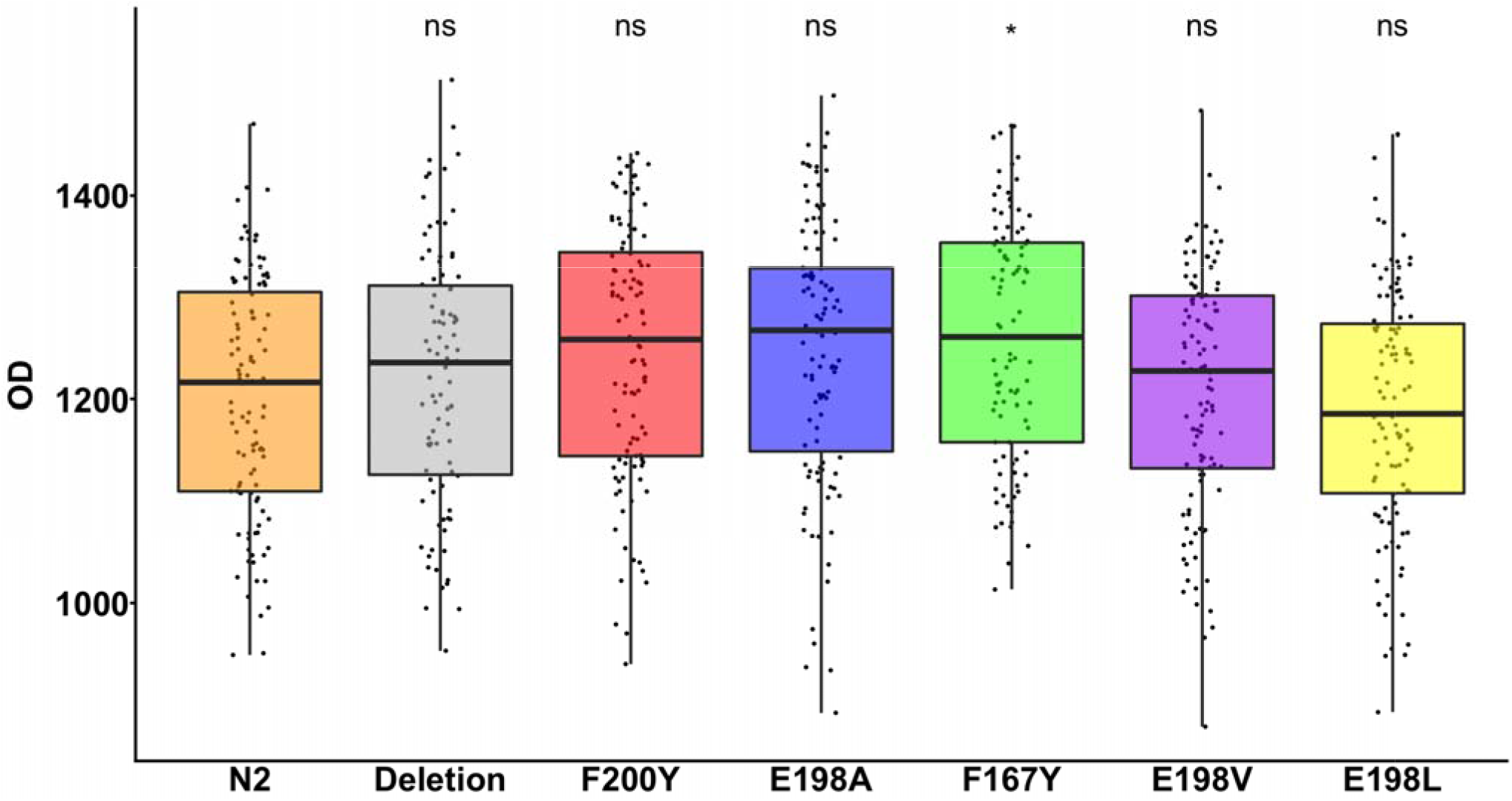
Strains containing parasite beta-tubulin alleles in control conditions. The x-axis indicates the allele introduced into the N2 strain. Median Optical Densities of populations with the indicated allele in control conditions (DMSO) are plotted on the y-axis. Each point represents a population of approximately 35 individuals. The F167Y strain is the only strain significantly different from the N2 strain in control conditions (p < 0.05, Tukey HSD).

**Supplemental Figure 3.**
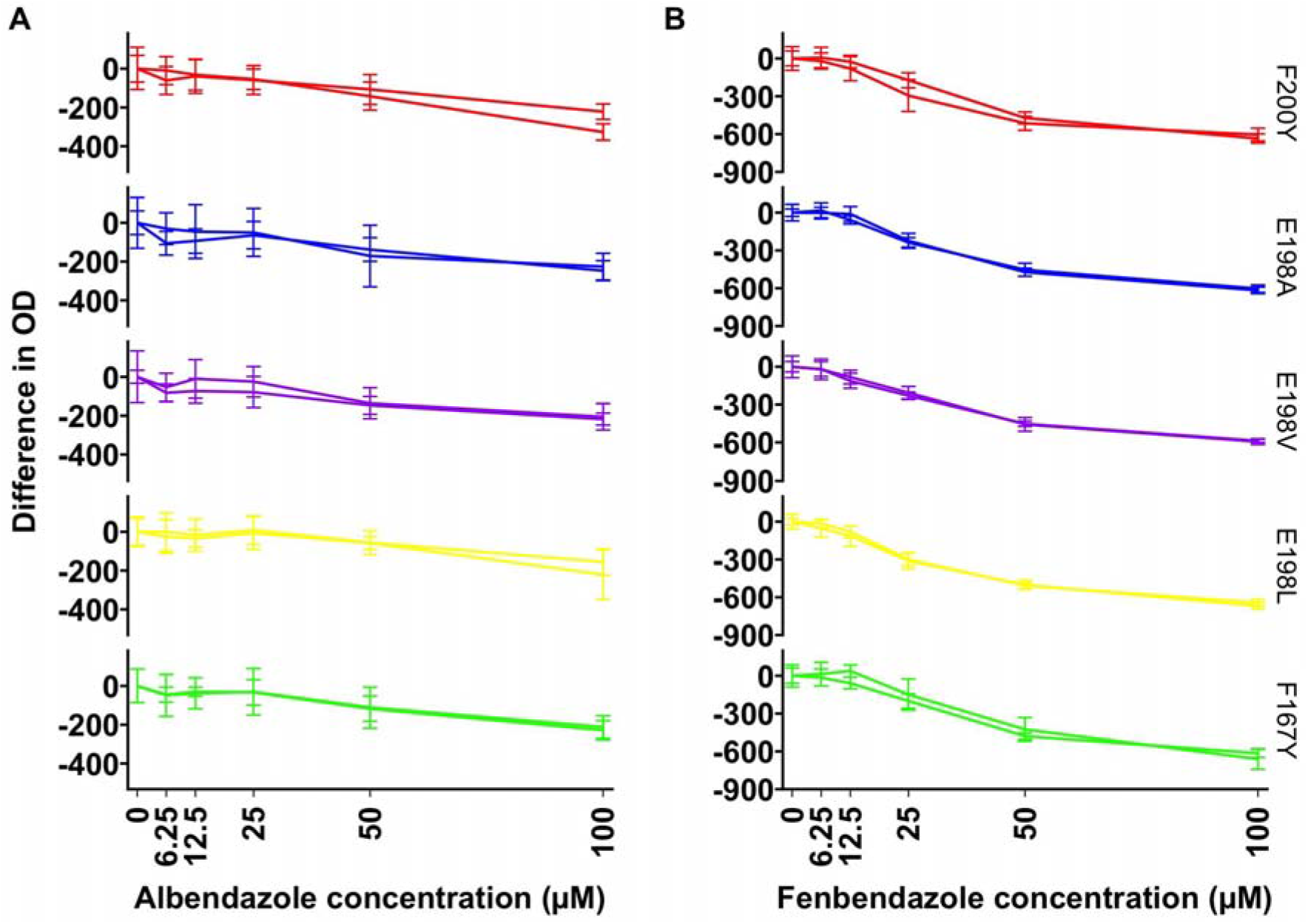
Dose-response assays with independently generated alleles. Drug-response assays of both canonical and non-canonical parasite beta-tubulin alleles in (A) albendazole and (B) fenbendazole. Normalized values were calculated by taking the phenotypic trait (median.EXT) value for each population and subtracting the mean median.EXT value of that strain at 0 μM. Normalized trait values are shown in responses to six drug concentrations (0, 6.25, 12.5, 25, 50, and 100 μM) of BZ. Data from two strains with the independent allele replacements are shown for each allele in both albendazole and fenbendazole. The allele replacement is shown to the right of (B) for each row.

**Supplemental Figure 4.**
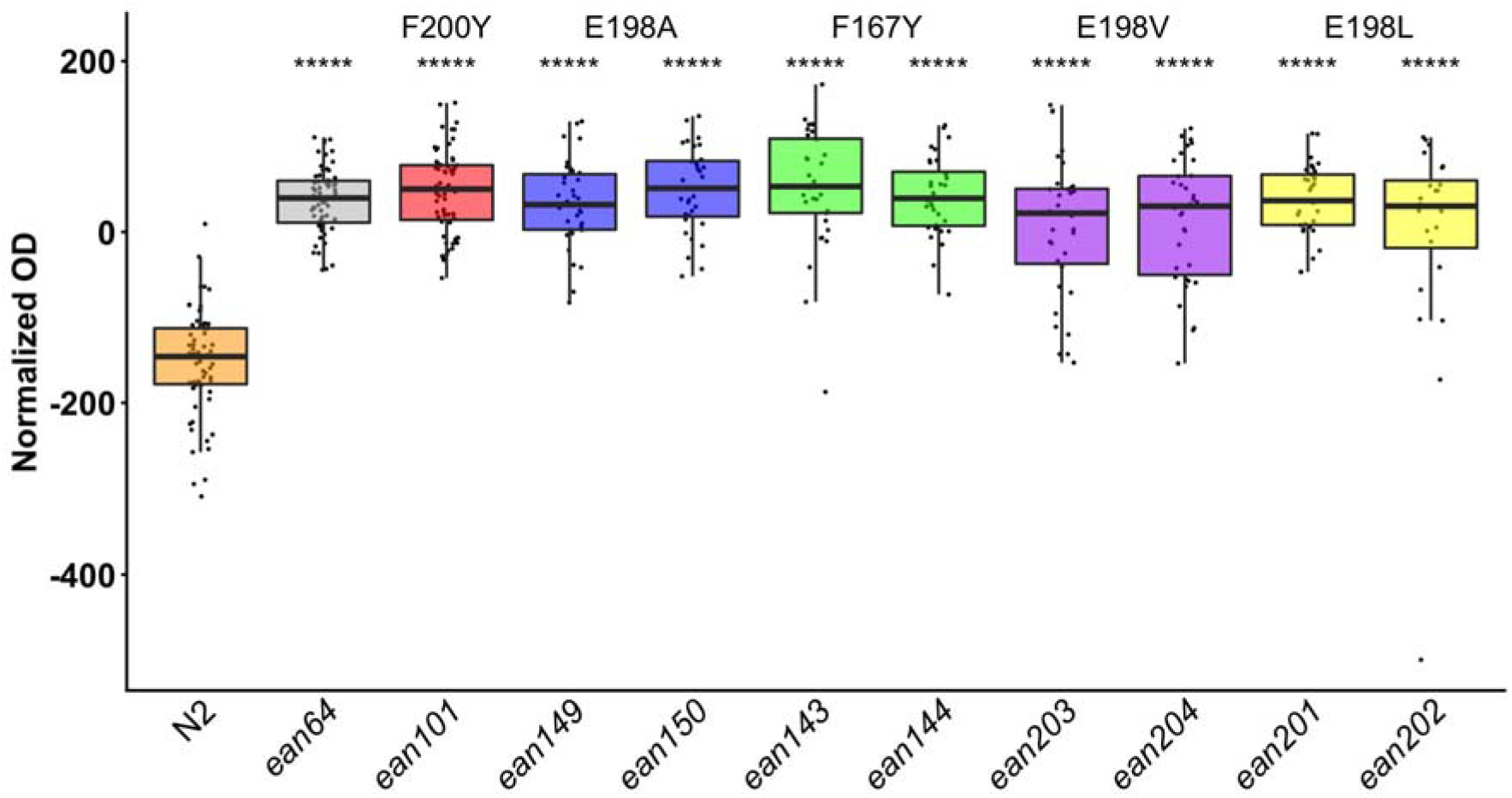
Independent alleles measured in the high-throughput assays at high replication. The x-axis denotes the *ben-1* allele designation of each tested strain (*ean64* = Δ*ben-1*, *ean101* = F200Y, *ean149* = E198A, *ean150* = E198A, *ean143* = F167Y, *ean144* = F167Y, *ean203* = E198V, *ean204* = E198V, *ean201* = E198L, and *ean202* = E198L). Regressed optical density of response to fenbendazole at 30 μM. Each point represents a population measurement of approximately 35-50 individuals. Independent edits of the same allele were not statistically different from one another (p-value < 0.0001, Tukey HSD).

**Supplemental Figure 5.**
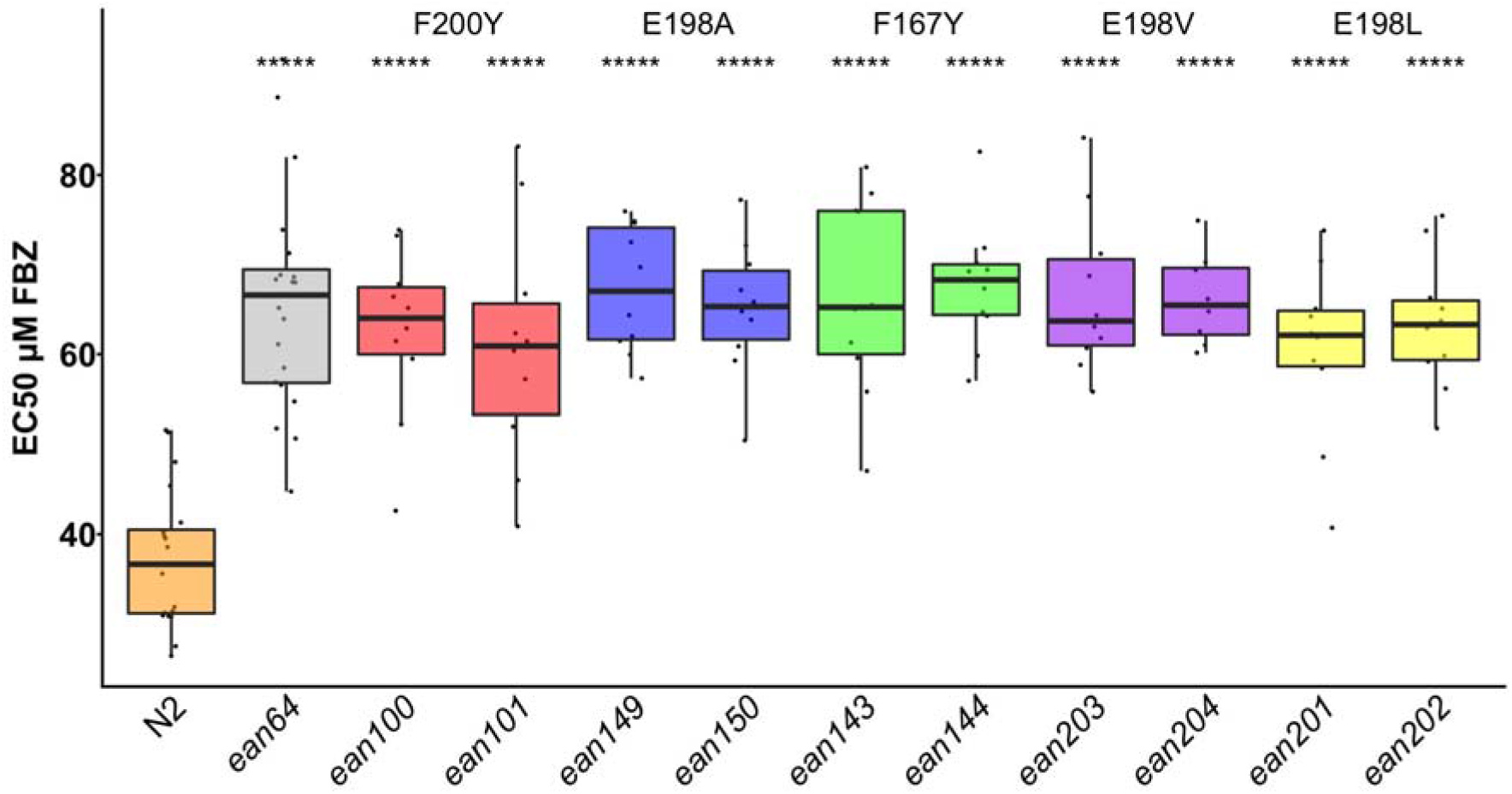
Estimated fenbendazole EC50 values for each the *ben-1* edited strains. The x-axis denotes the *ben-1* allele designation of each tested strain (*ean64* = Δ*ben-1*, *ean100* = F200Y, *ean101* = F200Y, *ean149* = E198A, *ean150* = E198A, *ean143* = F167Y, *ean144* = F167Y, *ean203* = E198V, *ean204* = E198V, *ean201* = E198L, and *ean202* = E198L). The y-axis shows the estimated EC50 in FBZ. Each point represents the estimated EC50 calculated from one replicate of our multi-dose drug assay. A replicate consists of one plate of the assay representing one well of each concentration with 35-50 individuals per well. Independent edits of the same allele were not statistically different from one another (p-value > 0.05, Tukey HSD). All alleles had significantly higher EC50 calculations than N2 (p-value < 0.0001, Tukey HSD)

